# Single cell kinetic modeling of redox-based drug metabolism in head and neck squamous cell carcinoma

**DOI:** 10.1101/2022.05.17.492281

**Authors:** Andrew D. Raddatz, Cristina Furdui, Erik Bey, Melissa L. Kemp

**Author notes:** **Corresponding Author:** Melissa L. Kemp.

## Abstract

Head and neck squamous cell carcinoma (HNSCC) cells are highly heterogeneous in their metabolism and typically experience elevated reactive oxygen species (ROS) levels in the tumor microenvironment. The tumor cells survive under these chronic oxidative conditions by upregulating antioxidant systems compared to healthy cells. Radiation and the majority of chemotherapies used clinically for treatment of HNSCC rely directly or indirectly upon the generation of short-lived ROS to induce cancer cell death. To investigate the heterogeneity of cellular responses to chemotherapeutic ROS generation in tumor and healthy tissue, we leveraged single cell RNA-sequencing (scRNA-seq) data to perform redox systems-level simulations of quinone-cycling β-lapachone treatment as a source of NQO1-dependent rapid superoxide and hydrogen peroxide (H_2_O_2_) production. Transcriptomic data from 10 HNSCC patient tumors was used to populate over 4000 single cell antioxidant enzymatic models. The simulations reflected significant systems-level differences between the redox states of healthy and cancer cells, demonstrating in some patient samples a targetable cancer cell population or in others statistically indistinguishable effects between non-malignant and malignant cells. Subsequent multivariate analyses between healthy and malignant cellular models point to distinct contributors of redox responses between these phenotypes. This model framework provides a mechanistic basis for explaining mixed outcomes of NQO1-bioactivatable therapeutics despite the tumor specificity of these drugs as defined by NQO1/catalase expression.

## Introduction

Head and neck squamous cell carcinoma (HNSCC) is one of the most prevalent types of cancer globally (1). Prophylactic measures such as HPV vaccination and the reduction of alcohol consumption and smoking are improving outcomes; however, five-year survival rates of HPV-negative HNSCC remain lower than 60% (2). While the etiology of HNSCC and anatomical locations within the oral cavity epithelial tissue are diverse, a hallmark of this cancer is elevated oxidative stress (3). Reactive oxygen species (ROS) at physiological concentrations are important as second messengers for many signaling processes including the MAPK, PI3K, NF-κB, and HIF pathways (4–8); however, ROS at higher levels promote tumorigenesis by causing genomic instability and proliferative signaling (9). If ROS levels are elevated even further, the level of oxidative stress cannot be managed and cells will go through one of several cell death mechanisms including necrosis, apoptosis, and ferroptosis (10,11). Cancer cells manage levels of ROS through multiple antioxidant enzyme systems (12), and under sustained oxidative stress will transcriptionally upregulate several antioxidant enzymes via the Keap1-Nrf2 axis (13,14). One treatment strategy is to selectively target cancer cells through the generation of reactive oxygen species (ROS) and disrupt the delicate balance these cells have between their higher antioxidant capacity and higher ROS levels (15–19). A unique approach to this strategy is utilizing NQO1-activatable quinone drugs to generate ROS. Because NQO1 is a quinone-reducing enzyme that is upregulated by Nrf2 (20), this approach should selectively target cancer cells that have constitutive Nrf2 activation. Furthermore, the generation of acute ROS by NQO1-activatible therapeutics causes a positive feedback response to more NQO1 expression, thereby enhancing the lethality of these compounds. Numerous studies have shown the benefit of these types of drugs alone, and targeting additional antioxidant and survival systems concurrently can improve the efficacy of the drug (21–24); however, there is debate as to whether the currently considered metric of NQO1:catalase expression or activity ratio is useful for identifying tumors susceptible to NQO1-activatable quinone drugs (25–28). To improve our understanding of the complex interplay between various antioxidant systems and the production of ROS by NQO1-activatable drugs, we developed and analyzed a differential equation model based on enzyme kinetic mechanisms that leverages the diversity of expression levels relevant to cancer redox systems. Furthermore, we explored potential uses for such a model by initializing parameter and species values using scRNA-seq data as a way to understand intratumor and patient variability in response to this type of chemotherapeutic intervention.

## Methods

### Cell Lines and Culture

HNSCC cell lines SCC-61 (Dr. Ralph R. Weichselbaum, The University of Chicago) and rSCC-61 (29) were cultured in RPMI-1640 cell culture media with L-glutamine (Caisson Labs, Cat#RPL03) with 10% FBS (Sigma-Aldrich, Cat#F4135) and 1% Pen/Strep (Caisson Labs, Cat#PSL01) at 37°C and 5% CO_2_. Cell media was changed every other day and cultures were passaged at 80% confluence and regularly tested for *Mycoplasma* (MycoAlert PLUS, Lonza, Cat#LT07).

### siRNA Transfection

3,000 cells were seeded in the wells of a black clear-, flat-bottom 96-well plate (Corning, Cat#3603). After 24 hours, cells were washed three times with PBS and siRNA packaged in lipid nanoparticles using the N-TER Nanoparticle siRNA Transfection System (Sigma-Aldrich, Cat#N2913) was applied to each well at 50 nM in 100 uL of serum-free media for 4 hours. After this, an equal volume of media with 2X FBS (20%) was added for the remaining 20 hours of transfection. For each gene, three of the top-performing predesigned MISSION siRNA constructs from Sigma-Aldrich were pooled and transfected concurrently. The transfection efficiency of these siRNAs against GAPDH has been performed previously (30) via Western Blot, and we repeated similar validation Western Blots with the pooled siRNAs against NQO1, a critical enzyme within our system (Figure S1). After 24 hours of transfection by siRNA, cells were washed three times with PBS and further experiments performed.

### β-Lapachone Treatment Response H_2_O_2_ Measurements

Following siRNA transfection and PBS washes, Amplex Red and Horseradish Peroxidase (HRP) (Thermo Fisher Scientific, Cat#A2188) were added to the wells and kinetic fluorescent reads of resorufin (excitation 571 nm, emission 585 nm), the product of Amplex Red and H_2_O_2_ in the presence of HRP, were taken to measure H_2_O_2_ over time. After 10 minutes of reads to determine baseline extracellular concentrations of H_2_O_2_, 3 μM of β-Lapachone (synthesized in Boothman Lab, Indiana University) was applied to cells in serum-free media and reads were taken for 2 hours.

### Ordinary Differential Equation Model Construction

The redox system ODE model was built upon a previously published model originally developed to describe the H_2_O_2_ clearance within Jurkat T cells in response to a bolus of extracellular H_2_O_2_ addition (12). The additional species included in new reactions are: oxidized extracellular β-lapachone (β-lap^ext^), intracellular O_2_•^-^, oxidized intracellular β-lapachone (β-lapQ), reduced intracellular β-lapachone (β-lapHQ), semioxidized intracellular β-lapachone (β-lapSQ), and glutathionylated intracellular β-lapachone (β-lap-GSH). New reaction rate terms are provided in Table 1. Supplemental Tables 1 and 2 list the complete parameters and initial values, respectively, used within the ODE system which were updated from the model originally characterized for Jurkat cells (12). ODEs were solved with ode15s in MATLAB R2020b, using a max timestep of 1 second.

**Table 1.**

| Reaction Name | Rate Term |
| --- | --- |
| β-lap permeation* | $k_{34} * A^{\text{cells}} * ([\beta\text{-lap}^{\text{ext}}] - [\beta\text{-lapQ}])$ |
| β-lap reduction | $k_{29} * [\beta\text{-lapQ}] * [\text{NADPH}]$ |
| β-lap semioxidation | $k_{30} * [\beta\text{-lapHQ}] * [\text{O}_2]$ |
| β-lap oxidation | $k_{31} * [\beta\text{-lapSQ}] * [\text{O}_2]$ |
| superoxide dismutase | $k_{32} * [\text{O}_2^{\bullet-}]^2$ |
| β-lap semireduction | $k_{33} * [\beta\text{-lapQ}] * [\text{NADPH}]$ |
| β-lap glutathionylation | $k_{35} * [\beta\text{-lapHQ}] * [\text{GSH}]$ |
| Glutathionylated β-lap permeation* | $k_{36} * A^{\text{cells}} * [\beta\text{-lapHQ-SG}]$ |

\*Permeation rate terms are divided by respective compartment volumes

### Sensitivity Analysis

Sensitivity values were calculated by increasing or decreasing parameter values by 10%, running the ODE solver for a simulated 2 hours, and using the following formula.

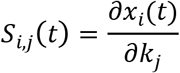

### scRNA-seq Data Analysis

HNSCC scRNA-seq data was collected from the gene expression omnibus (GEO Accession: GSE103322). This data had already been preprocessed to exclude cells with fewer than 2,000 genes detected or an average expression level below 2.5 of a curated list of housekeeping genes (31,32). Of the data from 18 patients, we retained the 10 patients that contained the most malignant cell transcriptomes as previously performed (31,33). t-SNE dimensional reduction was performed using the scikit-learn python library with default parameters besides PCA initialization. Enzyme abundance calculations from scRNA-seq data was performed as previously described (30). Briefly, kinetic rate constants from a mechanistic model of RNA production, RNA degradation, protein production, and protein degradation were used to determine equilibrium protein abundances given RNA levels. For proteins where these rate constants were not given, linear regression between RNA and protein was used to estimate protein abundance. Partial least squares regression (PLSR) was performed with log-transformed and zero-mean unit variance standardized data in SIMCA. Plots were generated using Seaborn and Matplotlib python libraries. The kernel density estimate plot was generated with default parameters using seaborn.kdeplot(). Scipy was used to conduct the Welch’s t tests with stats.ttest_ind() and equal_variance set to False.

## Results

### A systems level model of ROS generation by quinone cycling

We developed our model system to encompass three main aspects: 1) sets of critical H_2_O_2_-stabilizing antioxidant systems; 2) metabolism of the xenobiotic drug β-lapachone; and 3) the permeation of key species across membranes of the cell, including organelle-specific transport. We assumed that mitochondrial ROS production would remain constant due to basal respiratory metabolism, and mitochondrial antioxidant systems were not included, nor did we factor in activation of NADPH oxidases as a source of ROS as the consumption of NADPH by NQO1-catalyzed cycling of β-lapachone would render the NADPH oxidases inactive. Another assumption made was that due to high catalytic rates of NQO1 and antioxidant enzymes, 2 hours of simulated time was sufficient to capture the dynamics of the system. The relatively short period of simulated time allowed us to ignore transcriptional and translational regulation, such as how increased cellular oxidation would trigger Nrf2 nuclear translocation and upregulation of antioxidant genes including NQO1; therefore, total enzyme concentrations were assumed constant. The system and directionality of reactions and transport are shown in Figure 1.

**Figure 1.**
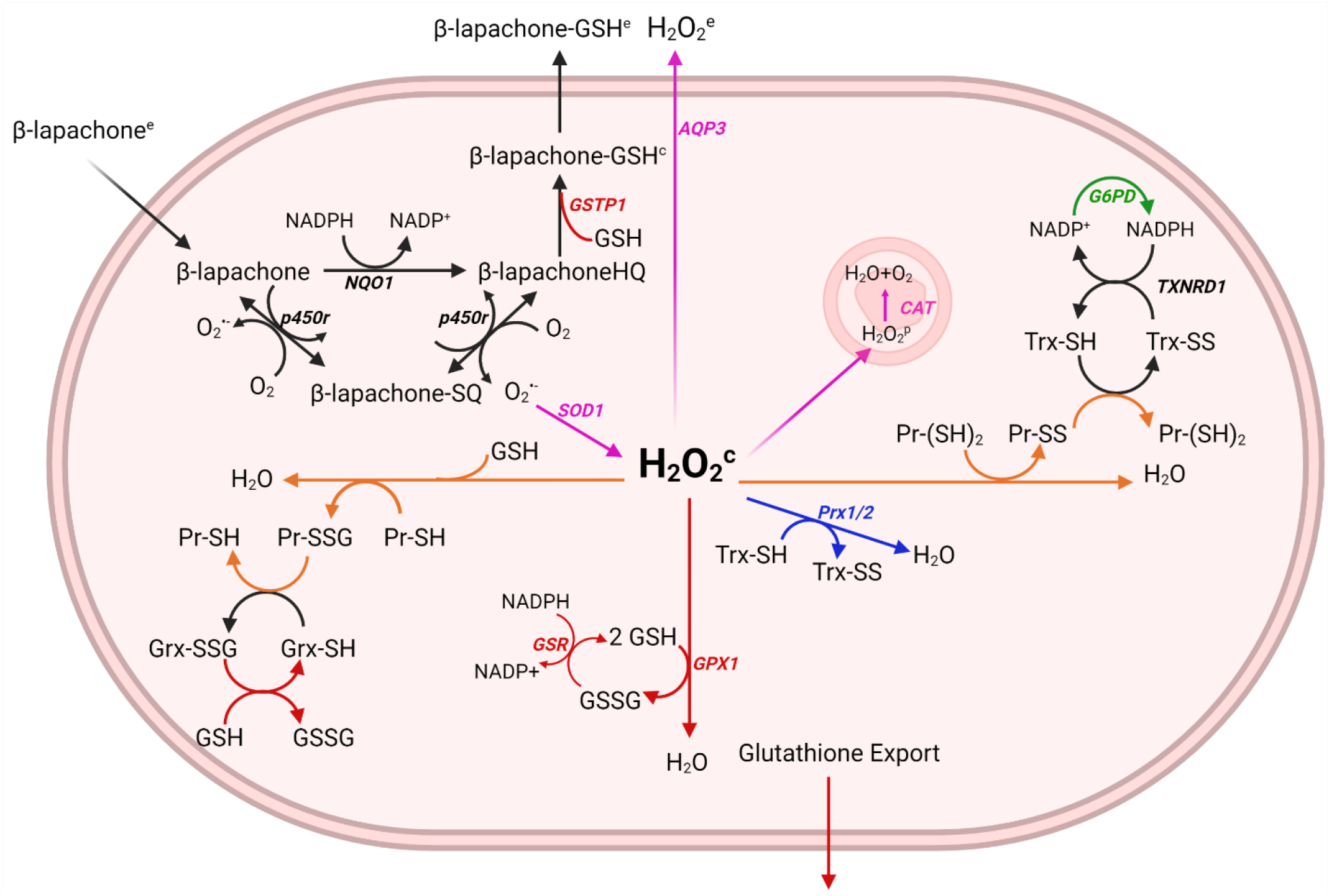
Generation of a relevant model of drug metabolism and hydrogen peroxide clearance pathways. The metabolism of β-lapachone by NQO1 results in the generation of superoxide (O_2_•^-^) and the oxidation of NADPH. Superoxide dismutase 1 (SOD1) in the cytosol converts the superoxide to hydrogen peroxide (H_2_O_2_) which is converted to water and oxygen by antioxidant systems including the peroxiredoxin/thioredoxin/thioredoxin reductase/sulfiredoxin system, the glutathione peroxidase/glutathione/glutathione reductase system, catalase, and the oxidation of free protein thiols. NADPH often serves as the reductant for cycling these antioxidant enzymes and it is used to reduce β-lapachone, thus canonical metabolic reactions involved in the production of NADPH are also included, such as glucose-6-phosphate-dehydrogenase (G6PD).

### Head and neck squamous cell carcinoma cells exhibit heterogeneity of redox gene expression

We sought to understand how variation in redox profiles of *in vivo* HNSCC tumors may reflect the distributed control of H_2_O_2_ clearance in tumor cells. To take advantage of new highly resolved omics technologies that provide rich tumor characterization, we analyzed scRNA-seq data from 10 HNSCC patients originally collected by Puram et al (31). In this dataset, there is a varying degree of cell type representation from each patient, likely due to both cross-patient tumor microenvironment heterogeneity and preprocessing of scRNA-seq reads for quality control. After splitting the dataset into malignant and non-malignant cells and reducing the variables to just 35 redox genes represented in our quinone cycling systems model, t-SNE clustering revealed malignant cells tend to cluster by patient (Figure 2a), suggesting that there are distinct, patient-based tumor redox profiles. We observed after clustering that tumors across patients have overall similar NQO1 levels, but that individual tumors display heterogeneity with respect to the distribution of cells expressing higher NQO1 (Figure 2b). This heterogeneity can also be observed when inspecting TXNRD1 and GLUD1 expression (Figures 2c, 2d). With this knowledge of heterogeneity between and within patient tumors, we leveraged redox transcriptional profiles per cell per patient to explore potential ROS buildup on cell- and tumor-based scales.

**Figure 2.**
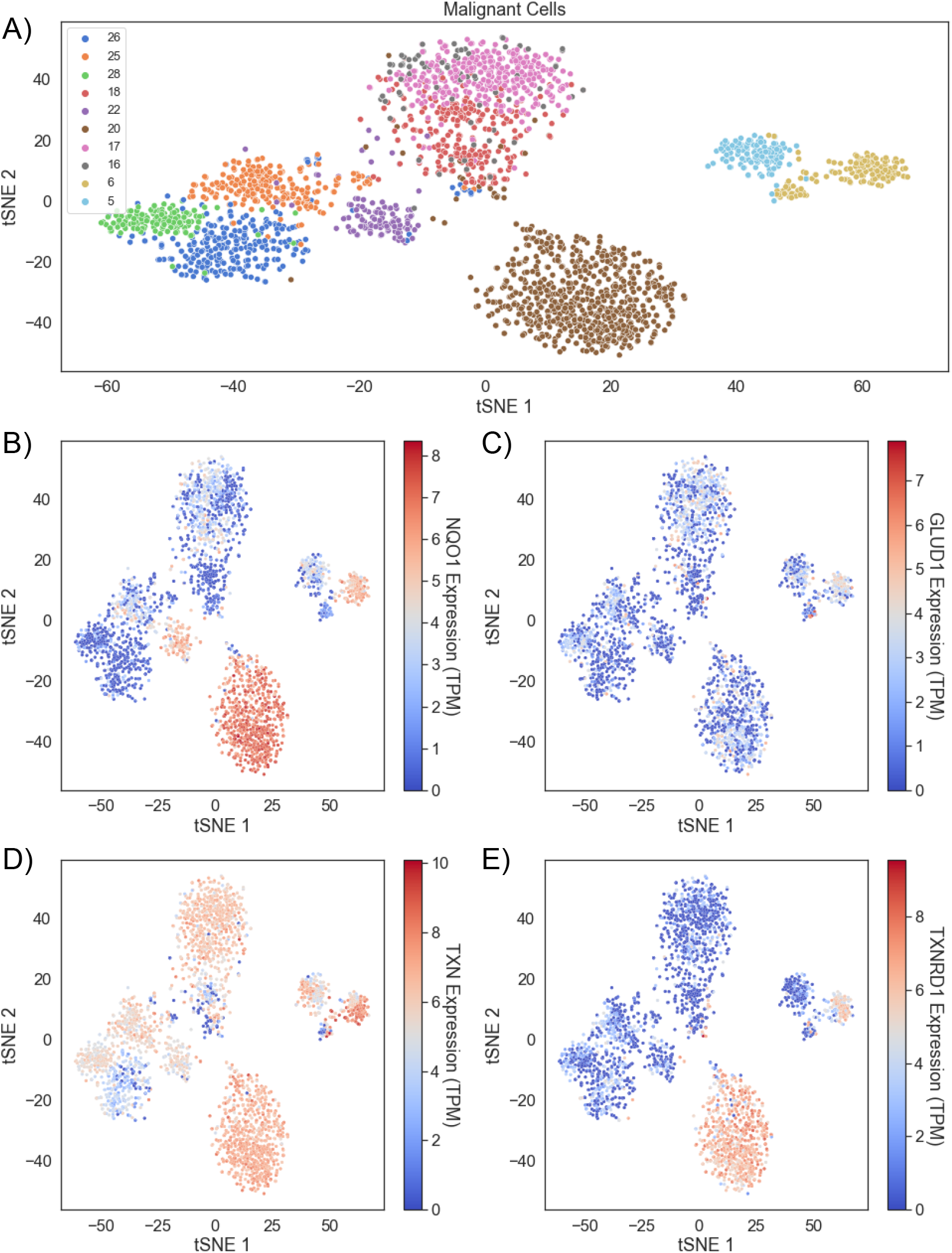
Head and neck cancers demonstrate intratumor and intertumor redox heterogeneity. Head and neck cancers demonstrate intratumor and intertumor redox heterogeneity. A) Malignant cells from 10 HNSCC patients cluster together based on redox profiles. B) Clusters colored by NQO1, C) GLUD1, D) TXN, and E) TXNRD1 expression.

### Initializing single cell ODE models with scRNA-seq

While scRNA-seq data has been widely used for exploratory data analysis and to understand gene expression correlations within developing tissues and cancer, this form of characterization has only recently been used to inform mechanistic kinetic models (34). We generated unique cell-based ODE systems using the previously analyzed scRNA-seq data. With the redox transcriptional profiles of almost 5000 cells from 10 patients, we first estimated the redox protein profiles as previously described (30,35) and imported these protein concentrations and related rate constants into our ODE model followed by simulation of the redox metabolism for each cell undergoing acute ROS generation by β-lapachone treatment. Specifically, AQP3, GSR, TXNRD1, NQO1, SOD1, POR, G6PD, and GLUD1 expression levels were used to adjust reaction rate constants by multiplying the rate constants by the percent change in the single cell expression from the average. G6PD and GLUD1 both generate NADPH and were combined into a single reaction in the model. GPX1, CAT, PRX1, PRX2, TXN, and GLRX expression levels were used to estimate initial enzyme abundances. PRX1 and PRX2 expression levels were combined and represent a single reaction in the model. All other parameters and species levels were kept from prior modeling (12).

### Sensitivity analysis shows H_2_O_2_ production is insensitive to individual enzymatic parameters

After constructing the ODE system, we sought to understand how influential each simulation parameter was on our system by performing a sensitivity analysis. We assessed the effect on intracellular H_2_O_2_ as the output variable of interest by altering model parameters up or down 10%. With sensitivities remaining below 1 and H_2_O_2_ only being somewhat sensitive to several parameters, we concluded that no single parameter could alter the H_2_O_2_ production significantly (Figure 3a).

**Figure 3.**
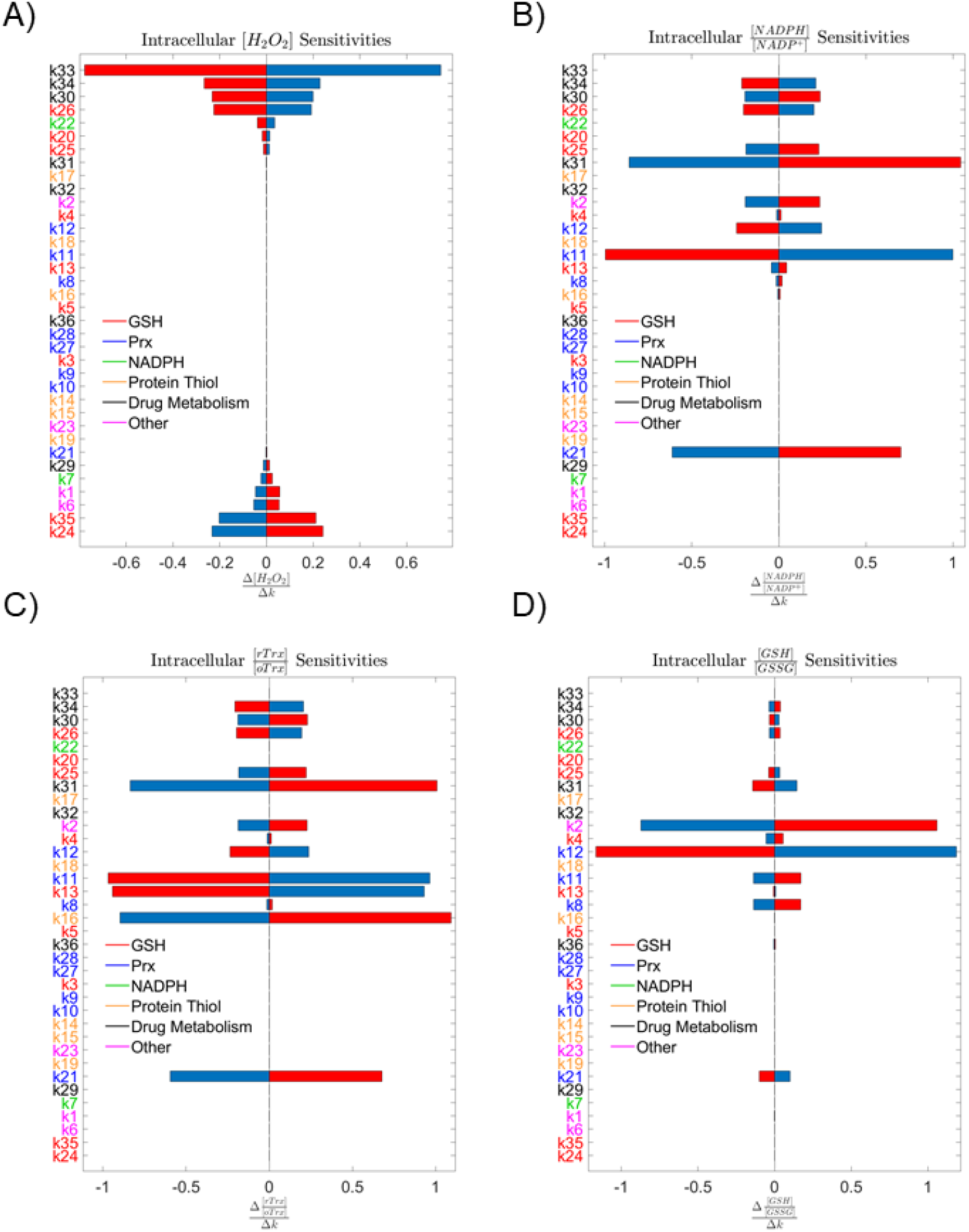
Sensitivity analyses. A) Analysis of system sensitivity to single parameter 10% perturbations colored by antioxidant subsystem shows low sensitivity of A) intracellular H_2_O_2_, B) NADPH:NADP+, C) Trx-SH:Trx-SS, and D) GSH:GSSG at 2 hours to any single parameter.

Parameter labels colored by antioxidant subsystem also indicate no single antioxidant system is controlling a majority of the H_2_O_2_ scavenging load. Expanding the number of outcomes to include redox ratios of reduced glutathione to oxidized glutathione, reduced thioredoxin to oxidized thioredoxin, and NADPH to NADP^+^ allowed us to assess the impact of these parameters on alternative indicators of redox status within the cell. The distribution of parameter importance in the sensitivity analyses across multiple redox mechanisms suggests that the reductive capacity of a cell is robust, and no single antioxidant enzyme system is predominantly responsible for clearance of H_2_O_2_ (Figure 3b-d).

### Experimental knockdown of antioxidant enzymes confirms model sensitivities

To experimentally validate the model, we used siRNA to perturb antioxidant enzyme levels and then observed the knockdown effect on acute H_2_O_2_ production induced by β-lapachone over a 100 minute period (Figure 4a). We confirmed via NQO1 Western blots that 24 hours of siRNA exposure leads to approximately 50% knockdown of expressed protein (Figure S1). We also probed NQO1 after silencing Nrf2 and PRDX1 and observed changes in NQO1 expression (Figure S1), suggesting either a global siRNA impact on ROS-related protein expression as demonstrated by Kippner et al. (36) or an indirect, downstream effect of these proteins on overall redox state. One possibility is that lowering Nrf2 and PRDX1 can increase basal ROS levels resulting in the production of antioxidant enzymes such as NQO1 through other mechanisms. After confirming pooled siRNA silencing, we knocked down a set of antioxidant or antioxidant-related enzymes including CAT, GPX1, GPX4, SOD1, GSR, PRDX1, TXN, TXNRD1, GLUD1, and G6PD to explore their impact on H_2_O_2_ production during β-lapachone treatment. We hypothesized that knockdown of antioxidants would result in an increase in H_2_O_2_, while knockdown of NQO1 would reduce drug metabolism and therefore H_2_O_2_ levels with β-lapachone treatment would be lower. We used Amplex Red to probe extracellular H_2_O_2_ levels over the course of 2 hours of drug treatment and compared the fold change in H_2_O_2_ relative to control scrambled siRNA (Figure 4b,c). With the maximum fold change no greater than 30% and relatively large standard deviation, these single knockdowns did not impact the redox state of the cells significantly by Welch’s t test, confirming our simulated parameter sensitivities.

**Figure 4.**
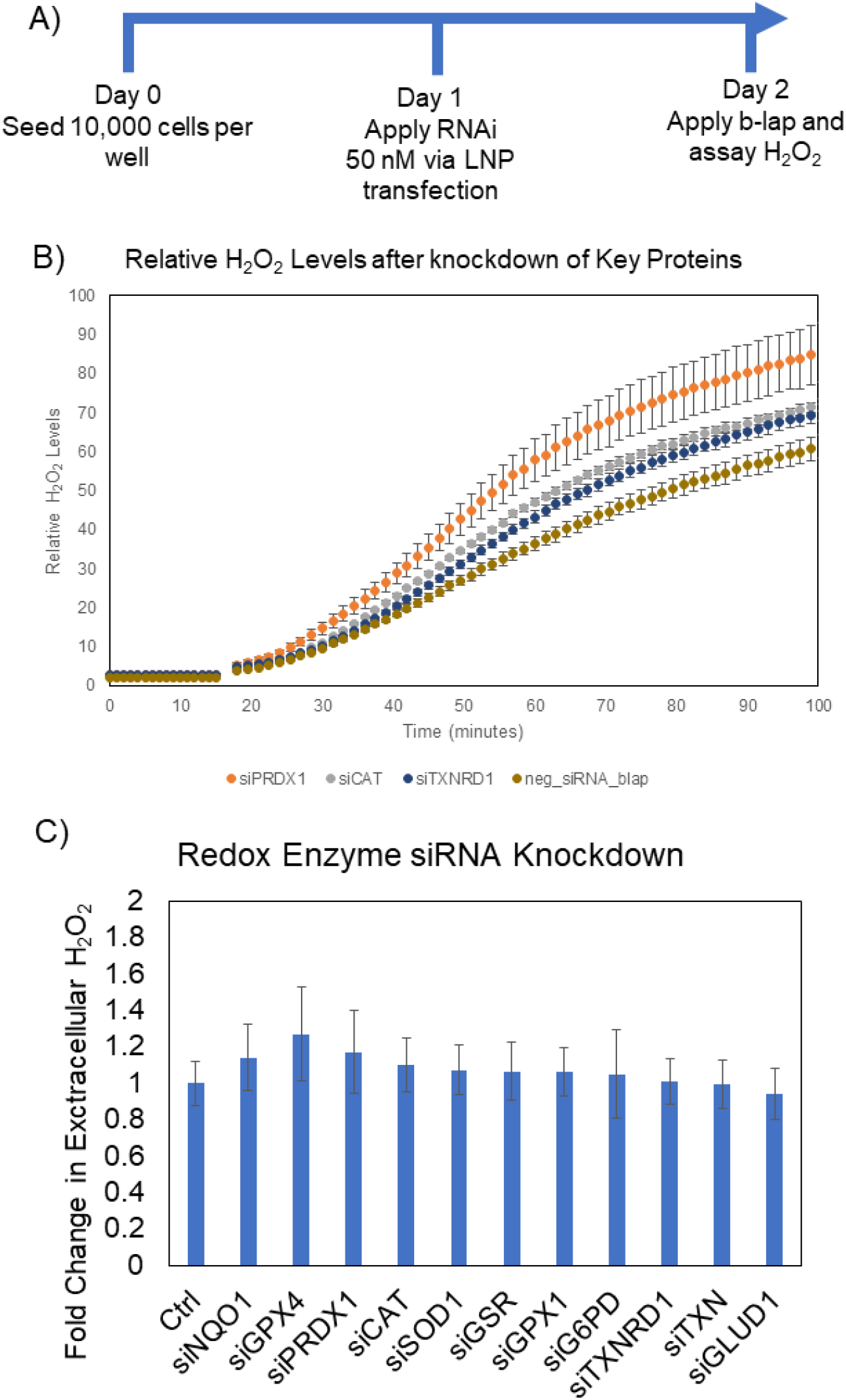
siRNA perturbation studies validate sensitivity analyses. A) siRNA knockdown workflow. B) Kinetic reads of H_2_O_2_ for PRDX1, CAT, and TXNRD1 enzyme knockdowns in a representative experiment showing increase in H_2_O_2_ after antioxidant knockdown (mean +/- s.d.). C) Aggregated fold change in H_2_O_2_ at 2 hours for each antioxidant knockdown shows limited increase in H_2_O_2_ confirming computational sensitivity analyses (mean +/- s.d).

### Comparison of H_2_O_2_ accumulation in healthy and cancer cells identifies patients with greatest potential for targeted therapy

Using this new system of generating single cell ODE models, the redox profiles of individual cells within HNSCC can vary greatly and result in a range of H_2_O_2_ spanning many orders of magnitude. After removing simulations that were unstable, we had 4,260 single cell simulation outputs across all ten patients. All of the ten patients showed a trend of more H_2_O_2_ generated by the malignant cells relative to the normal cells, with six patients exhibiting a statistical difference (Figure 5a).

**Figure 5.**
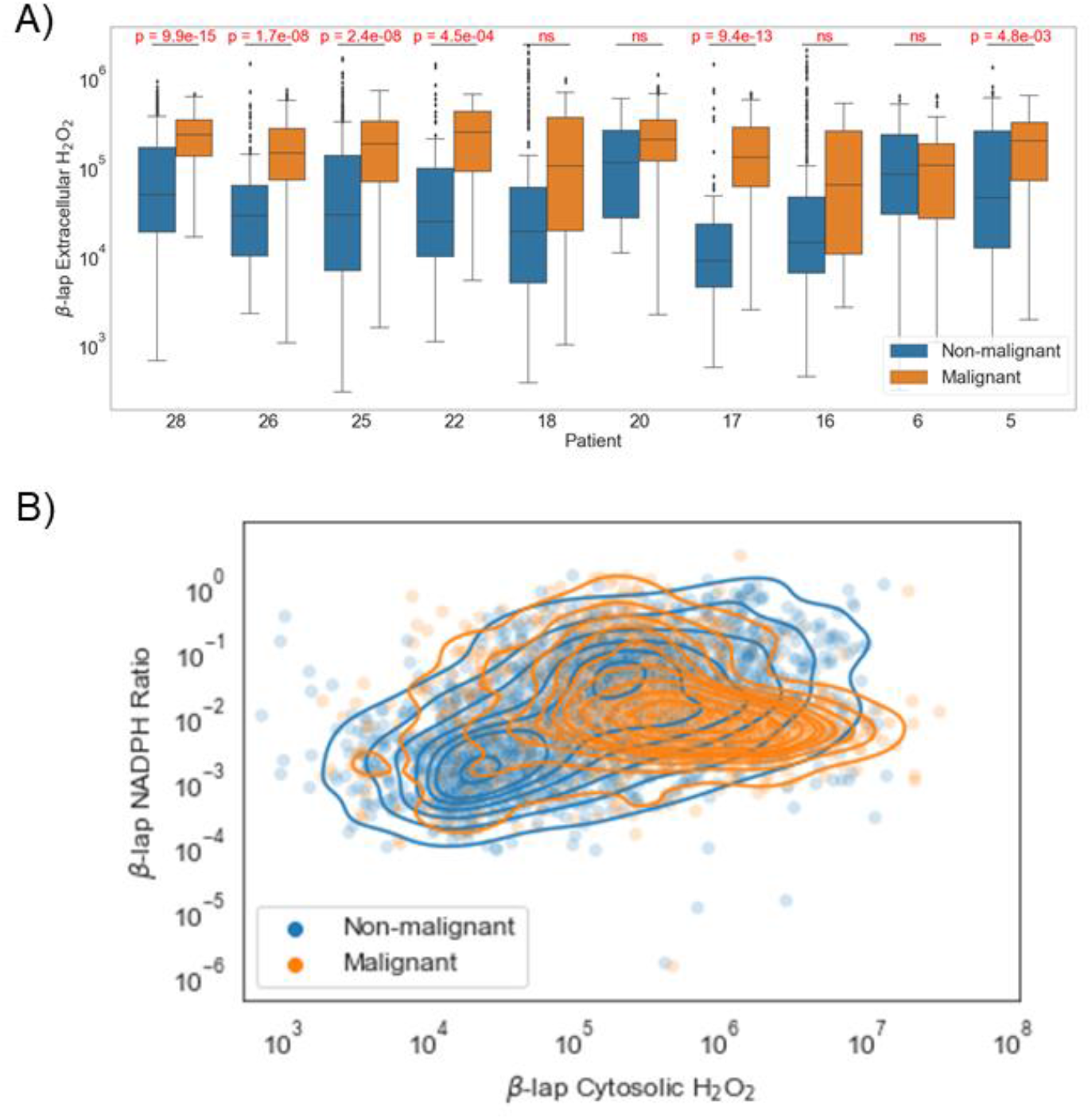
Model results using single cell gene expression values. A) Differences between extracellular H_2_O_2_ in healthy and malignant cells under β–lapachone by patient. B) Differences between NADPH:NADP+ ratio and extracellular H_2_O_2_ in healthy and malignant cells under β–lapachone.

Additionally, when comparing both H_2_O_2_ output and endpoint NADPH:NADP+ ratios across the 4,260 cellular models, we generally see higher H_2_O_2_ levels in cancer cells but no trend in NADPH:NADP+ ratios (Figure 5b). This shift demonstrates a potential for using single cell profiling to select patients for treatment with this targeted chemotherapy based on their redox profile. For the four patients where treatment induced H_2_O_2_ in both healthy and malignant cells without a statistically significant difference, the therapy may induce normal tissue toxicity impacting treatment and long-term quality of life.

### Initializing single cell ODE models with scRNA-seq identifies proteins correlated with H_2_O_2_ production

H_2_O_2_ concentrations and glutathione redox ratios after a 2 hour simulation were collected and used in partial least squares regression to probe the correlations between the protein concentrations within the model and the four output variables. With 7 and 6 components, respectively, both the malignant and non-malignant regression models are able to achieve both high explained output variance (non-malignant R^2^Y = 0.672, malignant R^2^Y = 0.689) and goodness of prediction (non-malignant Q^2^ = 0.656, malignant Q^2^ = 0.672). VIP scores identify NQO1, POR, TXNRD1, AQP3, and G6PD as the most important variables in the malignant model (Figure 6a) and POR, GLUD1, TXNRD1, NQO1, and GPX1 as the most important variables in the non-malignant model (Figure 6b).

**Figure 6.**
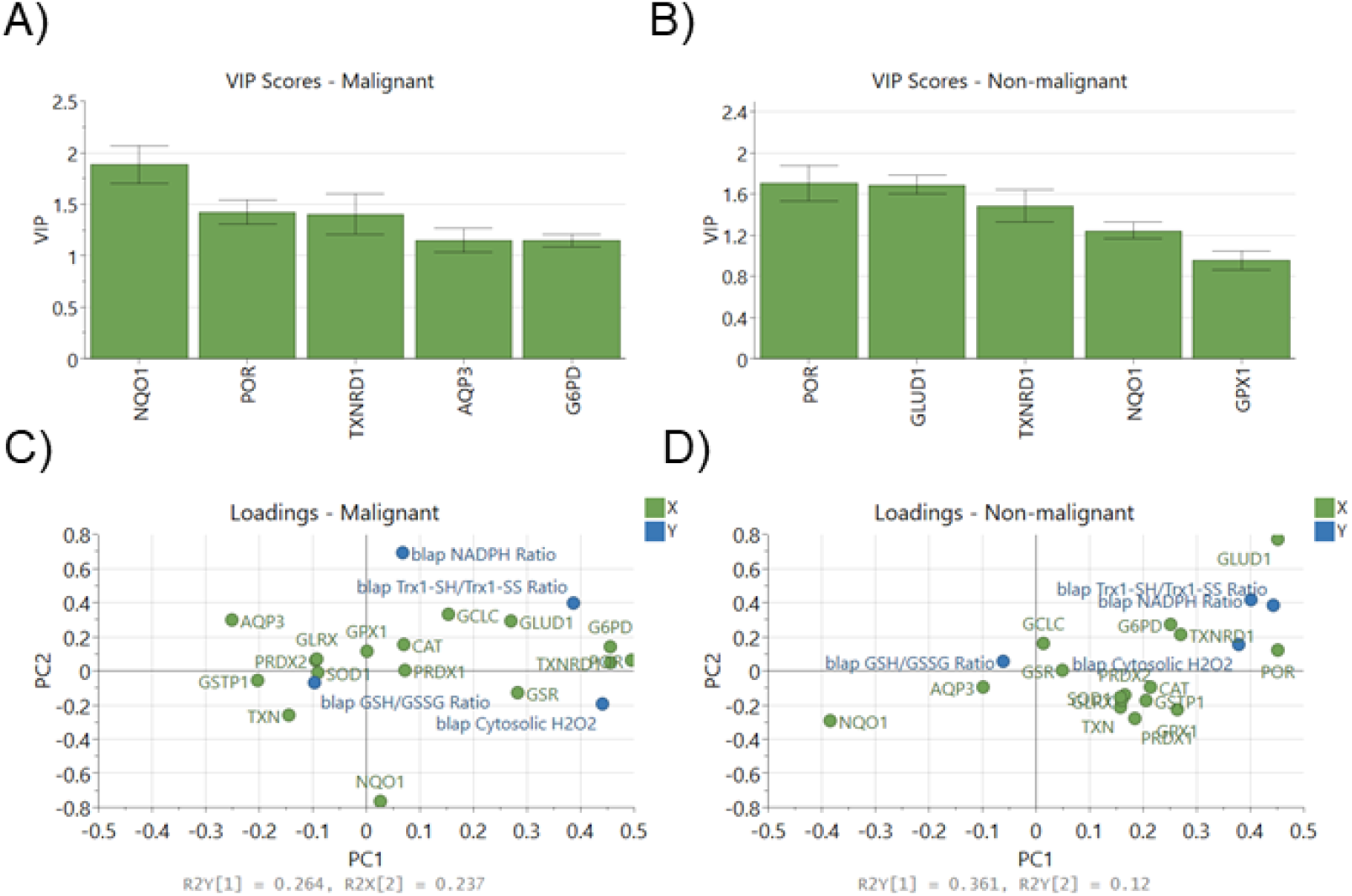
Partial least squares regression VIP scores and loadings. A) Genes with top 5 VIP scores in PLSR model in malignant simulations and B) non-malignant simulations. C) Breakdown of output into individual variables and loadings for each X and Y variable in malignant PLSR model and D) non-malignant PLSR model.

Collectively, the distribution of redox enzymes across principal components 1 and 2 differ between the two statistical models (Figure 6c,d). The distribution of NQO1 and CAT loadings in latent space reflects prior reports of NQO1/CAT as a useful metric in explaining the lack of β–lapachone lethality in non-cancerous tissues albeit not reflecting LD50 values across a diverse HNSCC panel (21). Additionally, the importance of POR demonstrates how the alternative path for β–lapachone reduction can still lead to significant superoxide production and therefore higher levels of H_2_O_2_. Expression of thioredoxin reductase 1 (TXNRD1) and NADPH-producing enzymes GLUD1 and G6PD are correlated with reduced thioredoxin levels, and most other antioxidant enzyme expression levels are less important due to low magnitude of their loading weights, i.e. proximity to the origin (Figure 6c,d).

## Discussion

Because the main mechanism of action by NQO1-activatable drugs is the generation of ROS, the ability for a cancer cell to manage ROS is a critical metric for chemotherapeutic response. NQO1:CAT ratio has been proposed as a predictive variable of NQO1-activatable drug success, but the utility of this metric is debated. Bey et al. in 2013 first speculated that NQO1:CAT could be useful after finding the use of exogenous catalase reduced the effects of β–lapachone in breast cancer (25), and higher NQO1:CAT were observed in NSCLC tumors that responded to treatment than in matched healthy tissue (26). In 2017 it was reported that the LD50 of β–lapachone did not correlate with NQO1:CAT in head and neck cancer (21). Additionally, while NQO1:CAT was not directly measured, inhibition of catalase and GSH did not lead to a sensitization of KEAP1-mutated NSCLC during β–lapachone treatment while inhibition of TXNRD and SOD1 sensitized cancers (28). A recent TCGA analysis revealed higher NQO1:CAT levels in hepatocellular carcinoma (HCC) than in matched healthy tissue, and the authors reported that the high NQO1 patient cohort had lower survival (37). These studies serve to highlight the complexity of the antioxidant system in the context of NQO1-activatable drugs like β–lapachone, and suggest the current approach for identifying how well a cancer would respond to the treatment is underdeveloped. In this report, we generated a more accurate model of ROS generation and scavenging under β–lapachone conditions by including additional antioxidant systems in an ODE-based approach in which H_2_O_2_ generation is a surrogate for drug potency. Including additional antioxidant systems and the kinetic information of enzymes simultaneously allowed us to predict measures other than NQO1:CAT that can serve as an indication of β–lapachone success.

When building a model to represent a biological system, there are always simplifications and assumptions that must be made using field expertise. Transcriptional regulation of the Keap1-Nrf2 axis on the scale of hours to days is not accounted for, in which the positive feedback of H_2_O_2_ activation of Nrf2-targeted genes results in enhanced NQO1 expression (14,38). Another major assumption used was that mitochondrial antioxidant systems would not reduce the large amount of ROS in this chemotherapeutic context due to the cytosolic location of NQO1 (39). Work done by Ma et al. shows that mitochondrial-targeted β-lapachone produces mitochondrial ROS using MitoSOX, while 3-hydroxy β-lapachone which is not mitochondrially targeted produces no substantial mitochondrial ROS (40).This allowed us to omit antioxidant enzymes expressed in the mitochondria such as SOD2, PRDX3, PRDX5. We did, however, find relatively high sensitivities of H_2_O_2_ permeabilities in the model, indicating the importance of how quickly a cell can export ROS during treatment. While H_2_O_2_ can passively diffuse through the phospholipid bilayer, it is also known to utilize aquaporin membrane proteins to travel through the plasma membrane (41–44). Because of the high sensitivities, measuring aquaporin expression levels could serve as a useful indicator of β-lapachone success.

When generating enzymatic models, direct expression levels of proteins can be acquired experimentally or from published datasets of other scientists’ experiments. We chose an alternative strategy by estimating protein abundance based on the scRNA-seq mRNA levels. Because transcriptional levels do not directly correlate to protein levels, we used a quantitative pipeline to estimate protein abundances that leverages previously published data from Schwanhausser et al (30,35). This allowed us to generate an ODE system specific to each cell sequenced in the scRNA-seq data. From our initial exploration of the scRNA-seq data, we observed the cells cluster by patient regardless of if they were healthy or cancerous similar to the results of an analysis conducted by Xiao et al. (33), so we concluded that each tumor was composed of a population of cells that were similar in redox profile. Yet when analyzing the expression of each antioxidant enzyme within these clusters, the overall antioxidant capacity or diversity of each tumor was unclear due to varied levels of each antioxidant enzyme. Our ODE model was able to stratify the patient tumors based on the differences in the expected response of healthy and cancerous cells to β-lapachone, shedding some light on the complex nature of redox systems. Because we used scRNA-seq data that had transcriptomes of both normal and cancerous cells, we were able to assess the relative dependence of these two cell populations on their antioxidant enzyme expression under oxidative stress. When the contours of the two cell populations are plotted in a 2D phase space of the two output variables, extracellular H_2_O_2_ and NADPH ratio, we find these overlap quite closely, but the cancer cell range is more compact. This suggests the cancer cell antioxidant phenotype can lead to a more controlled range of concentrations of ROS and reducing cofactors in oxidative environments, which can be seen as a survival advantage of the cancer cells. Similarly, while our comparisons of healthy and cancerous cells’ redox state after treatment were of the aggregated samples in each population per patient, there is wide variability within each group and some healthy cells show a more oxidatively stressed state than cancerous cells in the same tumor. The healthy cells represent a repertoire of components found in the tumor microenvironment ranging from fibroblasts to macrophages, and thus a diversity of responses to an oxidative insult is expected. While cancer cells are typically seen as being more oxidized, these results predict that tumor heterogeneity assessed at a single cell resolution can potentially challenge narratives established using bulk-based characterization.

A current issue with scRNA-seq data is a large volume of dropouts which leads to imputed values that are not true data (45). Methods for both higher quality sequencing and imputation are being developed, and as higher quality datasets are published this model can be updated to reflect that (46,47). Additionally, the added value of spatial information from new spatial omics technologies could further improve the model. With the model currently representing a single cell system, a multicellular model of all of the cells simultaneously with physical parameters included could better represent the tumor system and buildup and breakdown of ROS. Lastly, our model only predicts how these cells within patient samples would respond to β-lapachone. Working with directly validated samples is a more ideal workflow, and we look forward to testing these models’ accuracies if clinical data is made available in the future.

Altogether, this analysis demonstrates that developing a comprehensive enzymatic model of ROS generation and clearance using scRNA-seq data has the potential to identify the relative importance of various axes in the complex antioxidant network. We suggest that metrics other than NQO1:CAT should be considered when characterizing a HNSCC tumor and its capacity to respond to β–lapachone. These metrics include expression of TXNRD1, POR, and NADPH-producing enzymes such as G6PD and GLUD1. Ultimately, the systems approach outlined here demonstrates the value of utilizing mechanistic modeling in conjunction with omics data to attain a more comprehensive understanding of the cellular redox state.

## Notes

**Conflict of Interest Disclosure:** The authors declare no potential conflicts of interests.

### Competing Interest Statement

The authors have declared no competing interest.

