## Supplementary figures and images for "Single cell kinetic modeling of redox-based drug metabolism in head and neck squamous cell carcinoma"

### Supplemental Figures

#### Slide 1
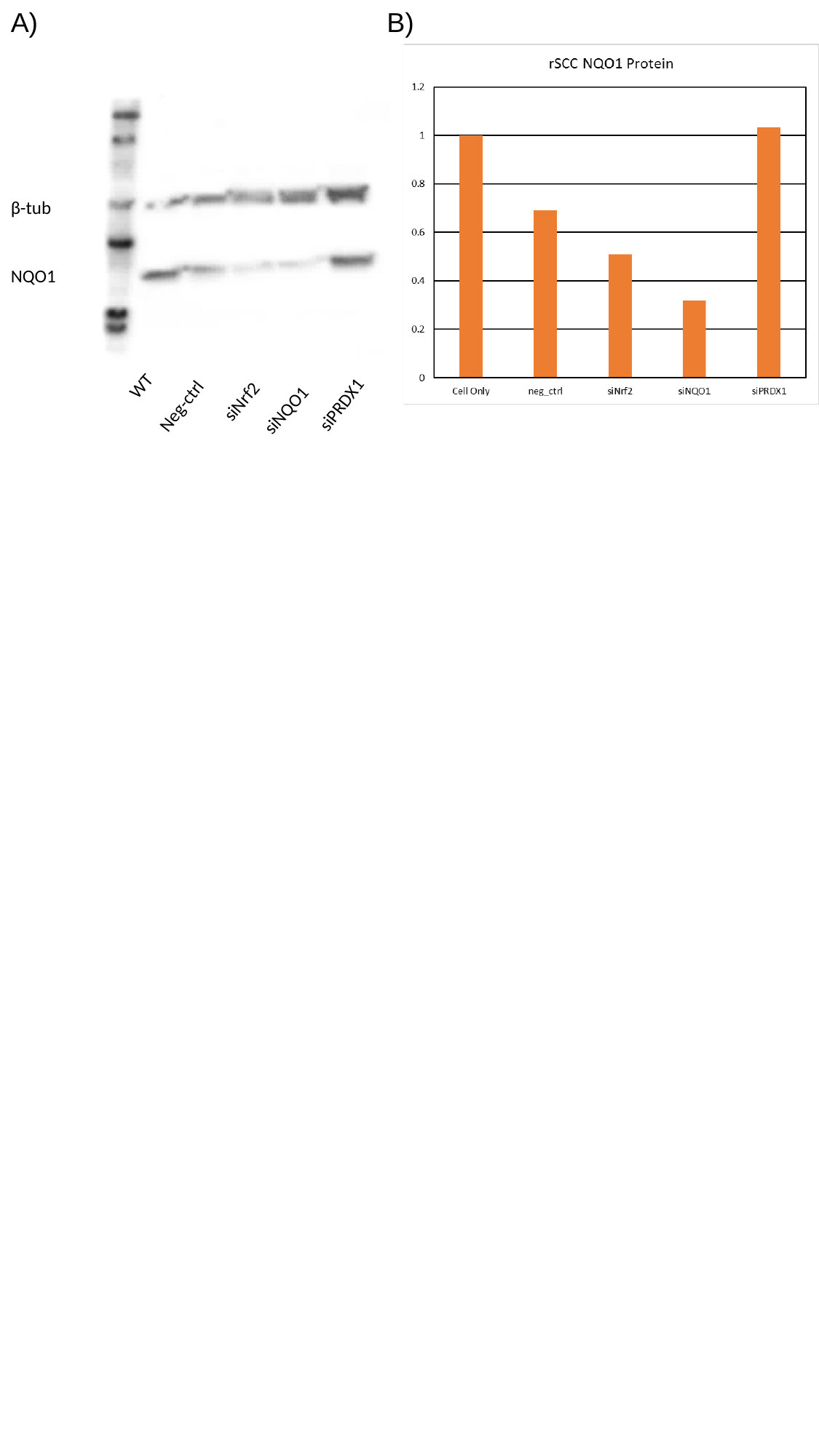

B)
A)
β-tub
NQO1
WT
Neg-ctrl
siNrf2
siPRDX1
siNQO1
